# Aerobiosis is not associated with GC content and G to T mutations are not the signature of oxidative stress in prokaryotic evolution

**DOI:** 10.1101/154534

**Authors:** Sidra Aslam, Xin-Ran Lan, Bo-Wen Zhang, Zheng-Lin Chen, Deng-Ke Niu

## Abstract

**Background:** Among the four bases, guanine is the most susceptible to damage from oxidative stress. Replication of DNA containing damaged guanines result in G to T mutations. Therefore, the mutations resulting from oxidative DNA damage are generally expected to predominantly consist of G to T (and C to A when the damaged guanine is not in the reference strand) and result in decreased GC content. However, the opposite pattern was reported 16 years ago in a study of prokaryotic genomes. Although that result has been widely cited and confirmed by nine later studies with similar methods, the omission of the effect of shared ancestry requires a re-examination of the reliability of the results.

**Results:** We retrieved 70 aerobe-anaerobe pairs of prokaryotes, and members of each pair were adjacent on the phylogenetic tree. Pairwise comparisons of either whole-genome GC content or the GC content at 4-fold degenerate sites of orthologous genes among these 70 pairs did not show significant differences between aerobes and anaerobes. The signature of guanine oxidation on GC content evolution has not been detected even after extensive controlling of other influencing factors. Furthermore, the anaerobes were not different from the aerobes in the rate of either G to T, C to A, or other directions of substitutions. The presence of the enzymes responsible for guanine oxidation in anaerobic prokaryotes provided additional evidence that guanine oxidation might be prevalent in anaerobic prokaryotes. In either aerobes or anaerobes, the rates of G:C to T:A mutations were not significantly higher than the reverse mutations.

**Conclusions:** The previous counterintuitive results on the relationship between oxygen requirement and GC content should be attributed to the methodological artefact resulting from phylogenetically non-independence among the analysed samples. Our results showed that aerobiosis does not increase or decrease GC content in evolution. Furthermore, our study challenged the widespread belief that abundant G:C to T:A transversions are the signature of oxidative stress in prokaryotic evolution.

## Background

Oxygen is an essential environmental factor for most organisms living on Earth, and its accumulation was the most significant change in the evolution of the biosphere and dramatically influenced the evolutionary trajectory of all exposed organisms [1]. Oxidative metabolism provides a large amount of energy to aerobic organisms and produces an unavoidable by-product: reactive oxygen species (ROS). ROS are highly reactive with most cellular organic molecules, including nucleotides and their polymerized products, DNA and RNA. Among the four bases, guanine has the lowest oxidation potential and is the most susceptible to oxidation [2]. The direct products of deoxyguanosine oxidation are 8-oxo-7,8-dihydro-guanosine (8-oxoG) and 2,6-diamino-4-hydroxy-5-formamidopyrimidine. As 8-oxoG has a lower oxidation potential than deoxyguanosine, 8-oxoG is susceptible to further oxidation into several hyper-oxidized products [3]. The replication of DNA containing these damaged deoxyguanosines can cause G to T mutations, the frequency of which depends on the efficiency of DNA repair enzymes and the accuracy of replication enzymes [3]. When the oxidatively damaged guanines are not in the reference strand, the mutations they caused would manifest as C to A mutations in the reference strand. Therefore, in some literatures the mutations resulting from oxidatively damaged guanines were denoted by G to T transversions while in other literatures they were denoted by G:C to T:A transversions. No matter which means of presentation, the G:C to T:A transversions were generally considered the hallmark of oxidative damage to DNA [4–7]. Consequently, oxidative DNA damage was generally believed to be a mutational force to decrease GC content [8–10]. Consistent with this idea, a negative association had been observed between metabolic rate and the GC content at the silent sites of animal mitochondrial genomes [11].

However, 16 years ago, Naya et al. [10] observed an entirely opposite pattern in which aerobic prokaryotes had higher GC contents than anaerobic prokaryotes in a comparison of whole-genome GC content using nonphylogenetically controlled statistics. Furthermore, these authors showed that the pattern was still evident when aerobes and anaerobes were compared within each major phylum of archaea and bacteria. Opposing to the widespread belief that oxidative stress causes frequent G:C to T:A transversions and decreases GC content, these results were described as “*counterintuitive*” [8]. The authors abandoned the neutralist interpretation to investigate possible selective forces, and they found that aerobes have lower frequencies of amino acids that are more susceptible to oxidation. As the non-synonymous sites of these amino acids are AT-rich, the high GC content of the aerobes might be explained by a deficiency of these amino acids. Moreover, they identified two potential benefits for aerobes with higher GC content. First, a high GC content might provide more stability to the DNA double strand, which would then be less accessible to oxygen radicals. Second, guanines located at synonymous sites might play a sacrificial role to protect other bases. This intriguing idea has been presented repeatedly [12, 13]. However, sacrificial guanine bases are easily mutated to T, and a mechanism is not available to maintain the sacrificial guanine bases during evolution [9]. Seven years later, the same group found that the GC content of microbial communities living in the dissolved oxygen minimum layer (770 m) is lower than that of communities living in other (either below or above) layers of the seawater column in the North Pacific Subtropical Gyre, thus emphasizing the link between aerobiosis and genomic GC content [14]. In contrast, three later studies on seawater columns ranging from tens to thousands of metres observed that the GC content of metagenomes tends to increase linearly with depth in marine habitats, with the lowest GC content observed in near-surface stratified waters [15–17]. Regardless of the data obtained for microbial communities inhabiting different seawater depths, the pattern of higher GC content in aerobes has been repeatedly observed in various nonphylogenetically controlled comparisons. Later studies by nine independent groups, each with their own criteria for selecting species, observed the same pattern [18–26].

A possible explanation of the counterintuitive observations is provided by artefacts resulting from the phylogenetic non-independence of the data [27]. In 2008 and 2010, two groups independently compared the whole-genome GC content of aerobes and anaerobes and accounted for the phylogenetic relationships [21, 28]; however, they did not find a significant association between aerobiosis and GC content in the prokaryotic species they studied. These findings have received very little attention, which was likely because the two publications did not focus on the insignificant relationship between aerobiosis and GC content. Since 2009, the study by Naya et al. [10] has been cited 86 times (Google Scholar; access date: May 15, 2018); however, only one of the cited studies explicitly noted the conflicting results: *“oxygen requirement [10] may (or may not [21]) have an impact on GC content”* [29]. The present study calls attention to these contradictory results. We took advantage of the rapid accumulation of sequenced genomes and performed an extensive investigation on the GC content and mutational spectrum in aerobic and anaerobic prokaryotes using a phylogenetically controlled method.

## Results and discussion

We first compared the genomic GC contents of the 1,040 aerobic samples and the 1,015 anaerobic samples without considering their positions in the phylogenetic tree. The genomic GC contents of the aerobic samples and the anaerobic samples are 56.46% ± 12.52% and 45.83% ± 11.03%, respectively. Two-tailed Mann-Whitney *U* test showed that the difference between them is highly significant (*P* = 5.6 × 10^−77^). Limiting this comparison within bacteria or archaea gave similar results (*P* = 5.6 × 10^−62^ and 1.8 × 10^−21^, respectively). Despite the much larger dataset, we also observed significantly higher GC content in aerobes than anaerobes. The reproducibility of this result is so high that the same pattern had been consistently observed in ten independent studies with nonphylogenetically controlled methods [10, 18–26].

To control the effects of a common ancestor, we performed a pairwise comparison between aerobes and anaerobes that are adjacent in the phylogenetic tree (Fig. 1). The difference in GC content within one pair is phylogenetically independent of the differences within any other pairs. Pairwise comparisons of the GC content between the selected aerobe-anaerobe pairs can thus be considered phylogenetically controlled comparisons. In this way, we did not find significant differences in the genomic GC content between aerobic prokaryotes and anaerobic prokaryotes (Fig. 2A, two-tailed Wilcoxon signed ranks test, *P* = 0. 826). When the pairwise comparison is limited to the 65 pairs of bacteria, the difference between aerobes and anaerobes remains statistically insignificant (two-tailed Wilcoxon signed-rank test, *P* = 0.883). Our phylogenetically independent comparison of genomic GC content gave a result that is different from the nonphylogenetically controlled comparisons [10, 18–26], but consistent with two previous studies that have accounted for the phylogenetic relationship [21, 28]. Still, we have not detected the signature of guanine oxidation.

**Fig. 1.**
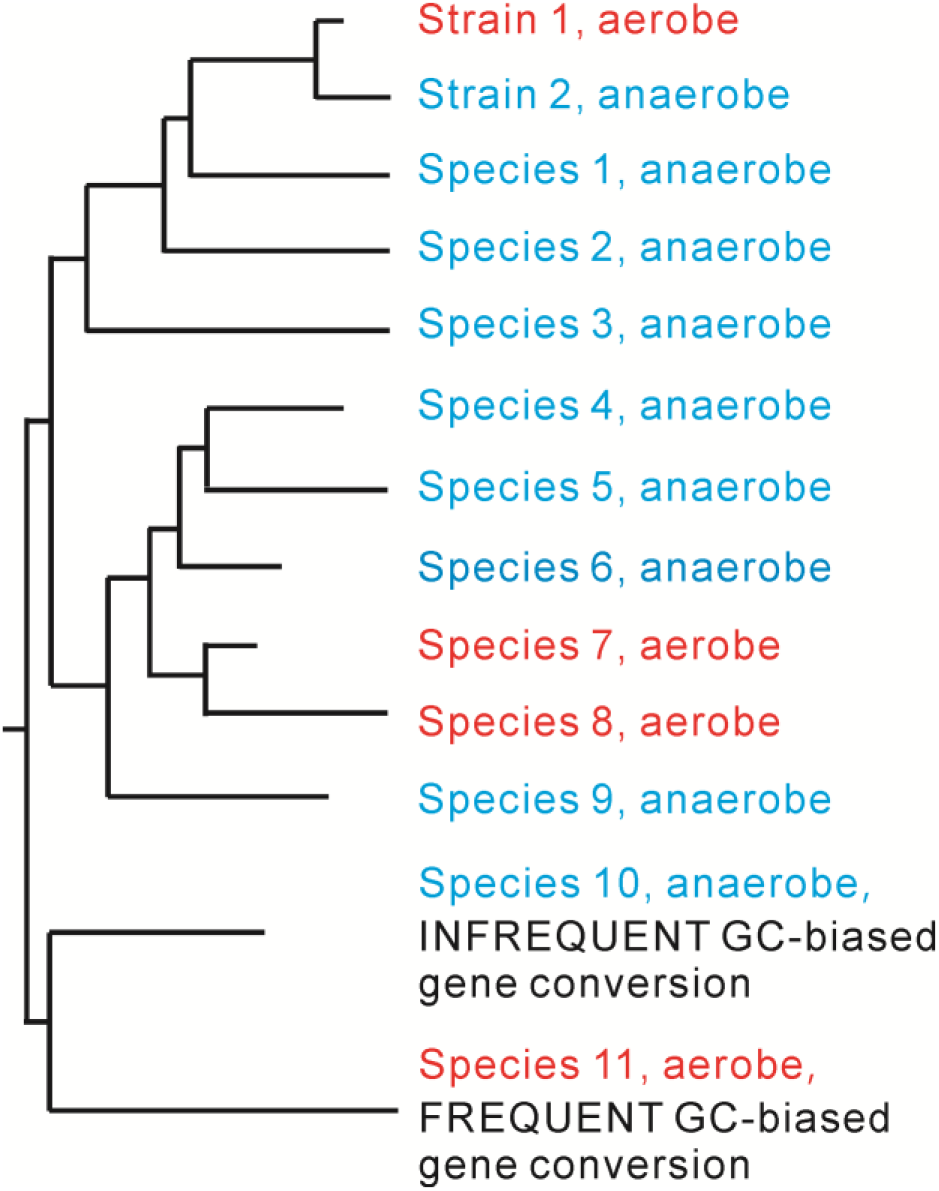
Illustration of the difference between nonphylogenetically-controlled comparisons and phylogenetically-controlled comparison performed in this study. In a nonphylogenetically controlled comparison, the aerobes (including strain 1, species 7, species 8, and species 11) are compared to all the anaerobes (including strain 2, species 1–6, and species 9–10). However, only three changes in oxygen requirement are observed in the illustrated evolutionary tree. The differences in GC content between these three branches are likely to be associated with changes in the oxygen requirement. Therefore, only three pairs should be included in a phylogenetically controlled comparison. For branches having multiple strains/species with different evolutionary rates (*e.g.*, species 4–8), we paired the slowly evolved aerobic strain/species with the slowly evolved anaerobic strain/species (species 6 *vs* species 7). In cases with two or more strains/species with identical divergence times, we preferentially selected the genomes in which more genes had been annotated. Next, the comparisons were duplicated using the dataset including the quickly evolved pairs (*e.g.*, species 5 *vs* species 8 selected from species 4–8). Nearly identical results were obtained in the duplicated comparison. The results of the former are presented in Fig. 2 and Table 1, and those of the latter are deposited as electronic supplementary material (Additional file 1: Fig. S1 and Table S1). The choice of an anaerobe from species 4, 5 or 6 or an aerobe from species 7 or 8 did not alter the results. The results of a pairwise comparison of the tips do not appear to be sensitive to the inaccuracies in the topology of the phylogeny.

Selective forces acting on non-synonymous sites might mask the specific effects of guanine oxidation within whole-genome sequences. For example, if codon GGG is mutated to TGG, this G to T mutation would be selected against because of the resulted change in the coded amino acid, from glycine to tryptophan. This exemplified mutation, even if occurs frequently, could not be fixed in evolution and so would not contribute to the evolution of GC content. In addition, the avoidance of oxidation-susceptible amino acids, of which the non-synonymous sites are AT-rich, might selectively increase the genomic GC content in aerobic prokaryotes [4]. The consequences of guanine oxidation, as a mutational bias, would be more accurately revealed by analysing the GC content of selectively neutral sequences or sequences under weak selection. Although the 4-fold degenerate sites (4FDS) might be under selection to maintain specific patterns of codon usage bias [52], they are by far the most common candidates for neutral or weakly selected sequences. Therefore, we performed pairwise comparison of the GC content at 4FDS. However, we did not find significant difference between aerobic prokaryotes and anaerobic prokaryotes (Fig. 2B, two-tailed Wilcoxon signed ranks test, *P* = 0.951).

**Fig. 2.**
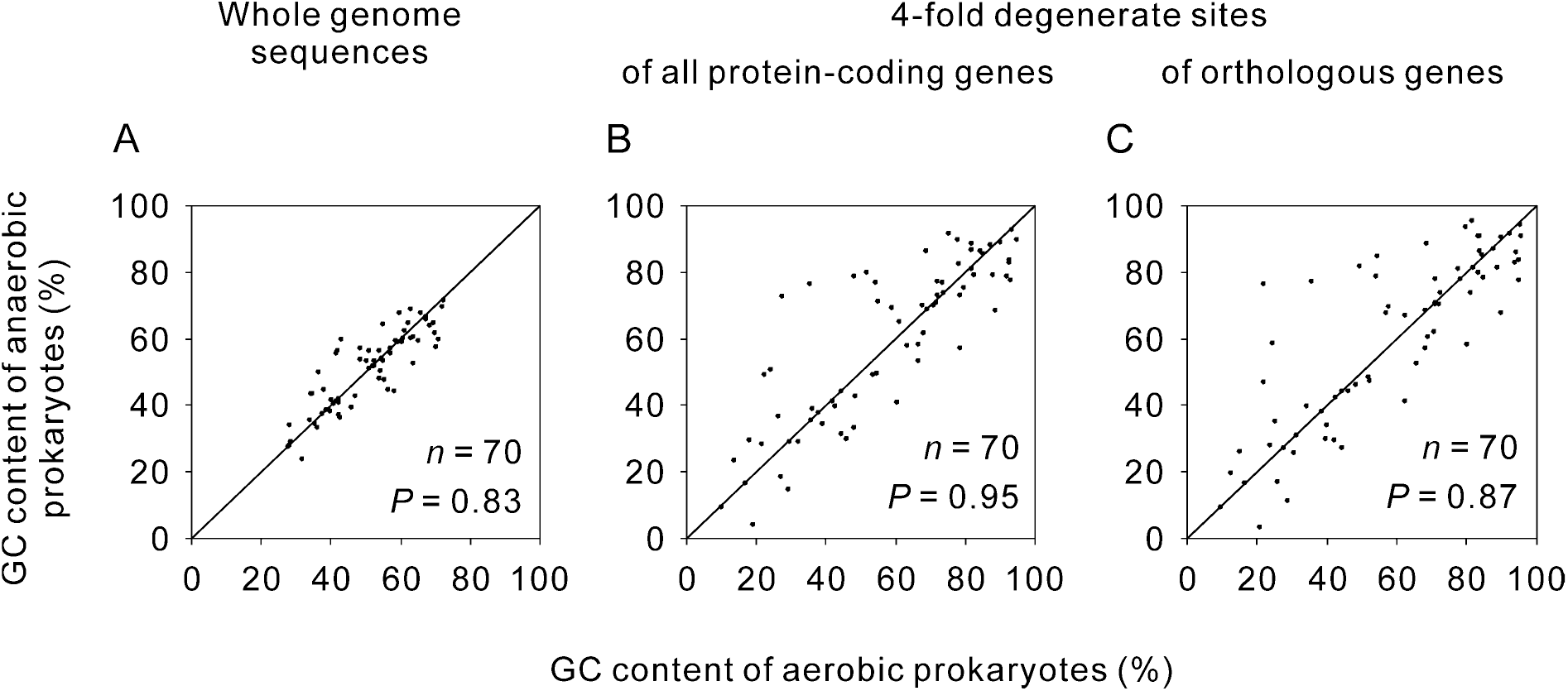
Pairwise comparison of GC content between aerobic and anaerobic prokaryotes. (*A*) Comparison of the GC content calculated from whole-genome sequences. (*B*) Comparison of GC content at the 4FDS of all protein-coding genes in each genome. (*C*) Comparison of GC content at the 4FDS of orthologous genes. The diagonal line represents cases in which aerobes and their paired anaerobes have the same GC content. Points above the line represent cases in which anaerobes have higher GC content than their paired aerobes, while points below the line indicate the reverse. All significance values were calculated using two-tailed Wilcoxon signed-rank tests.

Because horizontal gene transfer is extensive in prokaryotic evolution [60], the mutational force acting on the evolution of GC content in a lineage might be masked by the frequent horizontal transfer of DNA sequences with different GC content levels. The ideal genomic regions for comparison are sequences with orthologous relationships. For this reason, we compared the GC content of 4FDS within orthologous protein-coding genes. But still, we did not find significant difference between aerobic prokaryotes and anaerobic prokaryotes (Fig. 2C, two-tailed Wilcoxon signed ranks test, *P* = 0.886).

In addition to potential selective forces acting on non-synonymous sites and horizontal gene transfer, many other factors might increase the GC content of aerobes or decrease the GC content of anaerobes by specific mechanisms unrelated to changes in the oxygen requirement [8, 30]. GC-biased gene conversion has been widely observed as a driver of GC content increments [30, 31]. Organisms living at high temperatures tend to have higher GC contents in their structural RNA [32] and possibly in their whole-genome sequences (with debate, see [33–37]). G:C base pairs use more nitrogen and are energetically more costly than A:T base pairs; thus, AT-rich sequences may be favoured in non-nitrogen-fixing species and species living in challenging environments [8]. If guanine oxidation is a weak mutagenic force, then its effect on the evolution of GC content might be hidden by random combinations of these factors. Therefore, we propose that the relationship between oxygen requirement and GC content could be more accurately assessed if the oxygen requirement is the sole factor influencing the GC content that differs between each compared lineage. Although identifying all possible factors that influence the GC content of each species is impossible, distantly related species are more likely to differ in multiple factors that influence the GC content, whereas closely related aerobe-anaerobe pairs are more likely to differ only in the oxygen requirement, which is illustrated in Fig. 1. In addition to the oxygen requirement, species 10 and species 11 are assumed to differ in the frequency of GC-biased gene conversion. The frequent GC-biased gene conversion in species 11 might lead to a much greater increase in the GC content relative to the decrease in GC content caused by guanine oxidation. If so, aerobic species 11 would have a higher GC content than anaerobic species 10. Thus, we examined whether the relationship between oxygen requirement and GC content depends on the divergence time between the paired lineages. The divergence time between a pair of lineages was represented by the identity of their 16S rRNA molecules. We found that, no matter which threshold was used to define the close relatedness, the difference in GC content between closely related aerobes and anaerobes was not significant (two-tailed Wilcoxon signed ranks test, *P* > 0.10 for all the comparisons, Table 1 and Additional file 1: Table S1).

**Table 1.**
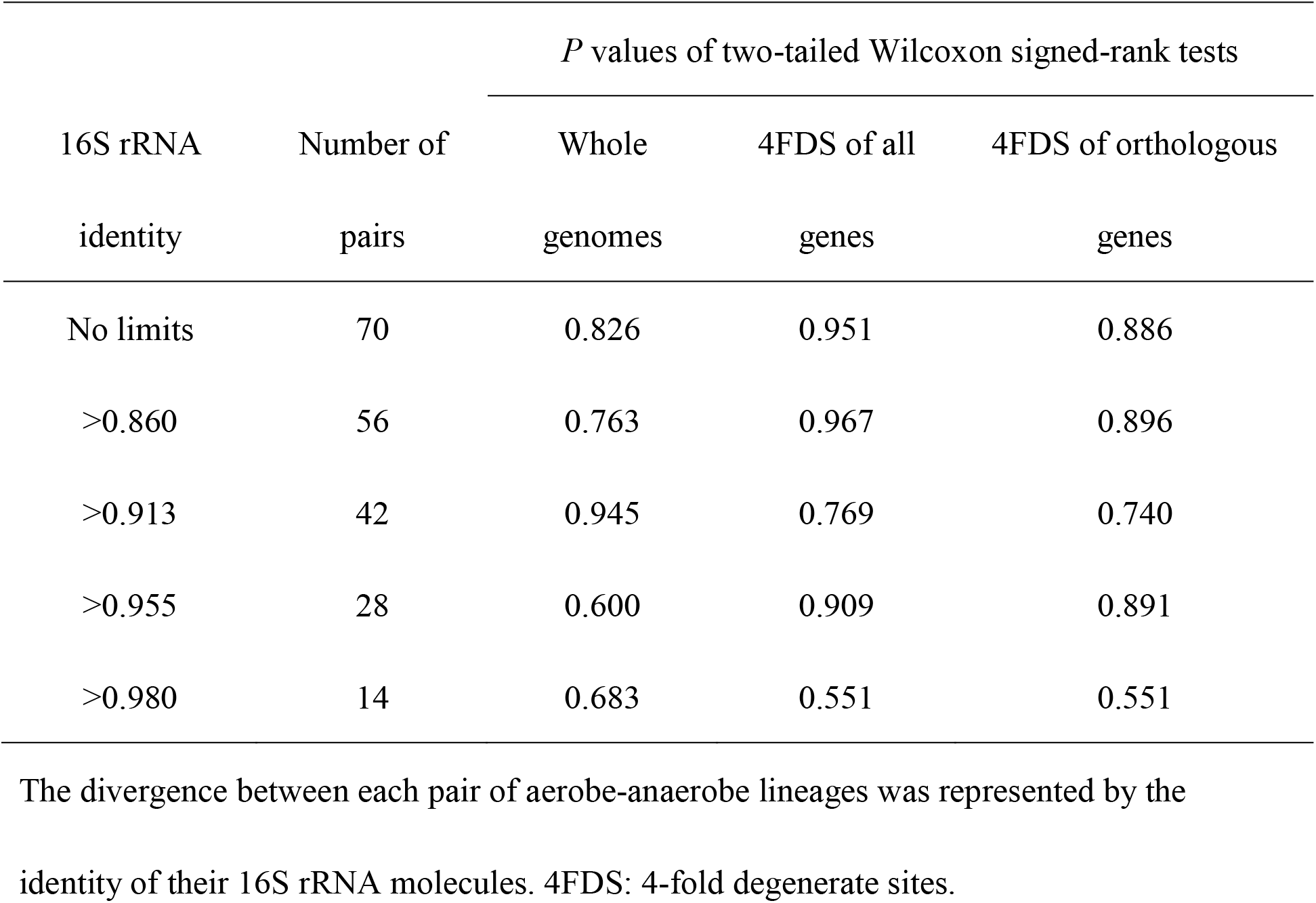
Relationship between GC content and aerobiosis is not dependent on divergence between compared lineages.

| 16S rRNA<br>identity | Number of<br>pairs | *P* values of two-tailed Wilcoxon signed-rank tests |  |  |
| --- | --- | --- | --- | --- |
|  |  | Whole<br>genomes | 4FDS of all<br>genes | 4FDS of orthologous<br>genes |
| No limits | 70 | 0.826 | 0.951 | 0.886 |
| >0.860 | 56 | 0.763 | 0.967 | 0.896 |
| >0.913 | 42 | 0.945 | 0.769 | 0.740 |
| >0.955 | 28 | 0.600 | 0.909 | 0.891 |
| >0.980 | 14 | 0.683 | 0.551 | 0.551 |

The divergence between each pair of aerobe-anaerobe lineages was represented by the identity of their 16S rRNA molecules. 4FDS: 4-fold degenerate sites.

In spite of elaborately controlling of other potential influencing factors, we did not detect any evidence for the signature of guanine oxidation on GC content evolution. One possible explanation is that efficient repair systems have been evolved and so the oxidative damage of guanine is only mildly mutagenic in most aerobic organisms [3]. If so, an anaerobe recently originated from aerobes would not have an obvious difference in GC content with its aerobic relatives. By contrast, an aerobe recently originated from anaerobes would need some time to evolve an efficient repair system. At this stage, frequent guanine oxidation would reduce the GC content. For this reason, we selected out eight orphan aerobes from our dataset. As the strain 1 illustrated in Fig. 1, the recent change in oxygen requirement of each orphan aerobe was supported by the existence of >3 close anaerobic relatives. We found eight orphan aerobes in our dataset. Pairwise comparison of these orphan aerobes with their anaerobic relatives did not show significant difference in either genomic GC content, GC content of 4FDS of all protein-coding genes, or GC content of 4FDS within orthologous genes (two-tailed Wilcoxon signed ranks test, *P* > 0.05 for all the three comparisons). As a control, we also compared the orphan anaerobes with their paired aerobes and did not find significant difference either (eight pairs, two-tailed Wilcoxon signed ranks test, *P* > 0.10 for all the three comparisons). Small samples cannot be ruled out as a source of the lack of significant differences.

After the above analyses, we could hardly reject the null hypothesis that aerobiosis is not associated with GC content in the evolution of prokaryotes. It is necessary to question the widespread belief that oxidative stress predominantly increases the rate of G:C to T:A mutations. The mutations retained in evolution may be inconsistent with that observed in experimental analyses [38]. We recalled two possibilities that were generally neglected. The first one is that anaerobic prokaryotes might also frequently undergo guanine oxidation. The antioxidant enzymes used by aerobes, like superoxide dismutase, have been identified in many obligate anaerobes [39–41]. The anaerobes might occasionally confront oxygen, and more likely suffer from the free radicals generated in oxygen-independent redox reactions and in radiolysis of intracellular water by ionizing radiation. Three enzymes, MutT, MutM and MutY, have well documented to be responsible for the repairing of oxidative damaged guanines [42]. Our preliminary survey showed that these enzymes are prevalent in anaerobic prokaryotes (Additional file 1: Table S2). Among the 70 anaerobic prokaryotes analysed in Fig. 2, genes encoding MutT, MutM and MutY have been detected in 41, 48, and 54 lineages, respectively. Meanwhile, in similar number of aerobic lineages (40, 51, and 57), the genes encoding these three enzymes have been detected. This result implicates the common occurrence of guanine oxidation in anaerobic prokaryotes. Consistently, the G to T and C to A substitution rates occurred at 4FDS in the orthologous genes of anaerobic prokaryotes were not lower than those of aerobic prokaryotes (two-tailed Wilcoxon signed ranks test, *P* > 0.60 for all comparisons). In addition, we did not detect any significant differences in the rates of the other 10 types of substitution (T to G, T to C, T to A, G to C, G to A, C to T, C to G, A to G, A to T, and A to C) or comparing only the orphan aerobes with the anaerobes they paired (Two-tailed Wilcoxon signed ranks test, *P* > 0.05 for all comparisons). According to the prevailing theory for mutation-rate evolution, natural selection tends to reduce mutation rates to the limit that is set by the power of random genetic drift [43]. The amount of oxidative damages left in aerobic genomes and anaerobic genomes after enzymatic repairing might depend on the power of random genetic drift, rather than the amount of mutagenic factors, like oxygen.

The second possibility is the existence of an opposite mutational force which cancelled the G to T mutation bias in aerobic prokaryotes. Replication of DNA whose guanines have been oxidatively damaged would result in G to T mutations. Meanwhile, guanine oxidation can also occur before incorporation of the guanine nucleotide into DNA [38, 42, 44]. During replication, 8-oxodGTP would be incorporated at the position of thymidine, pairing with adenosine. In the next round of replication, the 8-oxoG would be paired with cytidine if it happens to switch into the *anti* conformation. The resulted change is a T to G mutation. This type of mutation has been clearly revealed by mutant *E. coli* strain lacking the MutT enzyme [42], which is responsible for repairing oxidatively damaged dGTP. The two mutational forces, after being decreased in some proportions by the repairing systems, might cancel each other out in their effects on the evolution of GC content. In both the 70 aerobic prokaryotes and the 70 anaerobic prokaryotes analyzed in Fig. 2, the G to T transversion rates were a little higher than the T to G transversion rates. However, the differences were not statistically significant (Two-tailed Wilcoxon signed ranks test, *P* > 0.90 for both comparisons, Table 2). Surprisingly, the C to A transversion rates were not higher, but significantly lower than the A to C transversion rates in both aerobes and anaerobes (Two-tailed Wilcoxon signed ranks test, *P* < 0.05 for both comparisons, Table 2). This result does not support the generally expected higher frequency of G:C to T:A mutations resulting from the oxidative DNA damage associated with aerobiosis. Therefore, the G:C to T:A transversions should not be regarded as the signature of oxidative stress in prokaryotic evolution. We also compared other symmetric directions of mutations. No significant differences were observed between the rates of T to C and C to T or between A to G and G to A in aerobic prokaryotes (Two-tailed Wilcoxon signed ranks test, *P* > 0.05 for both comparisons, Table 2). However, different rates have been observed between all other pairs of symmetric mutational directions, A *vs*. T and C *vs*. G in the 70 aerobic prokaryotes and A *vs*. T, A *vs*. G, T *vs*. C, and C *vs*. G the 70 anaerobic prokaryotes (Two-tailed Wilcoxon signed ranks test, *P* < 0.05 for all comparisons, Table 2). Although these differences are unlikely associated with guanine oxidation or oxidative stress, they showed that there are some kinds of significant differences in our dataset and so indirectly support the validity of the observed insignificant differences.

**Table 2.** Comparison of the rates between symmetrical mutations in both aerobes and anaerobes.

|  | Aerobic prokaryotes |  | Anaerobic prokaryotes |  |
| --- | --- | --- | --- | --- |
| | Average ± SD | *P* | Average ± SD | *P* |
| G to T | 0.093 ± 0.085 | 0.917 | 0.094 ± 0.091 | 0.893 |
| T to G | 0.088 ± 0.069 | | 0.087 ± 0.072 | |
| C to A | 0.071 ± 0.073 | 0.001 | 0.063 ± 0.062 | < 0.001 |
| A to C | 0.146 ± 0.158 | | 0.134 ± 0.119 | |
| G to A | 0.085 ± 0.074 | 0.055 | 0.080 ± 0.074 | 0.009 |
| A to G | 0.122 ± 0.093 | | 0.124 ± 0.088 | |
| G to C | 0.131 ± 0.087 | < 0.001 | 0.129 ± 0.089 | < 0.001 |
| C to G | 0.096 ± 0.063 | | 0.098 ± 0.062 | |
| C to T | 0.129 ± 0.103 | 0.152 | 0.123 ± 0.109 | 0.025 |
| T to C | 0.170 ± 0.134 | | 0.174 ± 0.125 | |
| A to T | 0.095 ± 0.083 | < 0.001 | 0.094 ± 0.077 | < 0.001 |
| T to A | 0.061 ± 0.057 | | 0.065 ± 0.066 | |

All significance values were calculated using two-tailed Wilcoxon signed-rank tests. SD: standard deviation.

Although unexpectedly, these results are only new in prokaryotes. Similar results have been observed in recent spectrum analyses of somatic point mutations in mitochondrial DNA of aging tissues. A significant higher frequency of G to T mutations had not been detected in the mitochondrial genomes of aging animal tissues, being inconsistent with the contribution of oxidative stress to mitochondria-related aging [4, 45–47].

## Conclusions

Our phylogenetic independent comparison did not detect significant difference in GC content between aerobic prokaryotes and anaerobic prokaryotes. The result is different from nonphylogenetically controlled comparisons which always give a pattern of higher GC content in aerobes than anaerobes [10, 18–26]. Meanwhile, we did not detect significant difference in GC content even after elaborate controlling of other GC-content influencing factors. Our further analyses of the nucleotide substitution rates at 4FDS of orthologous genes showed that the mutations generally be attributed to guanine oxidation are not different in their frequency between aerobic prokaryotes and anaerobic prokaryotes. Moreover, guanine oxidation might exert two mutational forces simultaneously on the evolution of GC content evolution, both G to T mutations and T to G mutations. Different from the general expectation, our results indicated that aerobiosis is not associated the evolution of GC content in prokaryotes. Meanwhile, we suggested that the G:C to T:A transversions are not the appropriate signature of oxidative DNA damage in studies of prokaryotic evolution.

## Methods

In the Genomes Online Database (GOLD) [48], organisms are divided into ten categories according to their oxygen requirements: undefined, aerobe, anaerobe, facultative, facultative aerobe, facultative anaerobe, microaerophilic, microanaerobe, obligate aerobe, and obligate anaerobe. To avoid controversy, we retrieved only four categories: aerobe, anaerobe, obligate aerobe, and obligate anaerobe (access date: September 9, 2017). In the present study, the aerobes and obligate aerobes were merged into one group termed aerobes, and the anaerobes and obligate anaerobes were merged into another group termed anaerobes. In total, we obtained 4,009 aerobic prokaryotic samples and 2,707 anaerobic prokaryotic samples. The GC contents of 2,137 aerobic samples and 1,744 anaerobic samples were obtained from the summary section of the homepage of each species or strain in the NCBI Genome database. In the nonphylogenetically controlled comparison, we used the average value to represent the GC content of species that had multiple strains consistent in oxygen requirements. In species of both aerobic and anaerobic strains, the GC content of each strain was considered an independent sample. The genome sequences of the paired species or strains were retrieved from the NCBI Genome database (ftp://ftp.ncbi.nlm.nih.gov/genomes/). The GC content used in the pairwise comparison was calculated from the downloaded genome sequences rather than retrieved directly from the NCBI Genome database. Although the GC content values from these two sources were not identical, they were highly similar. The regression equation was y = 0.9927x + 0.0026, and the R^2^ value was 0.9909.

In the phylogenetically controlled comparison, we compared each aerobic prokaryote with its closest anaerobic relative. Therefore, we selected all the species that included both aerobic strains and anaerobic strains. Then, from the remaining species, we selected all the genera that included both aerobic species and anaerobic species. Finally, we selected the families that included both aerobic genera and anaerobic genera. Aerobic and anaerobic prokaryotes distributed in different families or higher taxonomic ranks were not included in our pairwise comparison. Referring to the All-Species Living Tree [49], we roughly filtered out the species that were unlikely to be usable for pairwise comparison of closely related aerobes and anaerobes. For example, in Fig. 1, species 1, 2, 3 and 9 were discarded during the rough filtration of the samples. For the remaining samples, we constructed a neighbour-joining tree using the p-distance model integrated in the software MEGA7 with 16S rRNA [50]. The p-distance (pairwise nucleotide distance) is the proportion of sites at which nucleotide sequences differ divided by the total number of nucleotides compared. The bootstrap values were obtained with 1,000 replications. For the poorly solved branches, we separately constructed their phylogenetic tree in the same way using 16S rRNA. In the four cases in which the phylogenetic relationships could not be resolved using 16S rRNA sequences, we constructed their phylogenetic trees using the *dnaj* gene sequence, which is another widely used phylogenetic marker [51–53]. Each difference in oxygen requirement between one pair of adjacent lineages was considered an event of evolutionary change in oxygen requirement (Fig. 1). The representative aerobic and anaerobic strains or species within each group were selected according to their branch lengths in the phylogenetic tree.

For a comparative analysis of the GC content at 4FDS in orthologous genes, we retained only the genomes whose protein-coding sequences had been annotated. In total, our dataset included 70 aerobe-anaerobe pairs.

For genomes in which the 16S rRNA gene annotations were not available, we identified the 16S rRNA genes by searching the genomes for the corresponding Rfam 13.0 profiles using Infernal (version 1.1.2) [54, 55].

We noticed that many bacterial genomes have not been fully assembled and some 16S rRNA sequences are fragmental. In the alignment of these 16S rRNA fragments, there are often large gaps not because of insertion/deletion occurred in evolution, but because of the incompleteness of the sequences. Both gaps and mismatches in the alignment are counted in the calculation of similarity, but only mismatches are counted in the calculation of identity. Identity is thus more solid than similarity in the comparison of fragmental 16S rRNA sequences. Therefore, we used the identity of 16S rRNA sequences to represent the divergence time between each pair of lineages. The sequences were aligned using ClustalW with its default parameters [56].

Orthologous genes between the paired lineages were first predicted by the reciprocal best blast hits and then screened using the program Ortholuge (version 0.8) using its default parameters [57, 58]. The thresholds of ratios 1 and 2 were both set to 0.8. Ortholuge is an ortholog-predicting method based on reciprocal best blast hits, and it improves the specificity of high-throughput orthologue predictions using an additional outgroup genome for reference. Ortholuge computes the phylogenetic distance ratios for each pair of orthologues that reflect the relative rate of divergence of the orthologues. Orthologues with a phylogenetic ratio that was significantly higher than that of the other orthologues in the genomes were considered incorrectly predicted and thus were discarded.

Properly aligned 4FDS of orthologous genes were obtained using the codon-preserved alignment software MACSE (version 1.2) with its default parameters [59]. Only the 4FDS that the nucleotide of one or both members of the aerobe-anaerobe pair were identical to that of outgroup were counted as the denominator in the calculation of the substitution rate. A substitution at a 4FDS was counted when the nucleotide of one member of the aerobe-anaerobe pair was different from that of outgroup while that of the other member was identical to that of outgroup.

Published sequences of MutY, MutM, MutT from the bacterium *Escherichia coli* str. K-12 substr. MG1655 (NCBI taxonomy ID: 511145) and the archaea *Azotobacter vinelandii* DJ (NCBI taxonomy ID: 322710) were used in bi-directional BLASTP [60]; database: non-redundant protein sequences; default parameters) to search the candidate homologous proteins in the respective pairs of bacteria and archaea, respectively.

## Additional files

Additional file 1: Fig. S1 and Table S1. Comparisons using the dataset containing the quickly evolved aerobe-anaerobe pairs. Table S2. Presence and absence of genes coding enzymes responsible 8-oxoG repairing in the aerobic and anaerobic genome studied in figure 2. (DOCX 251 kb)

Additional file 2: The data generated and analysed during this study. (ZIP 1585 kb)

## Abbreviations

ROS: reactive oxygen species; 8-oxoG: 8-oxo-7,8-dihydro-guanosine; 4FDS: 4-fold degenerate sites.

## Acknowledgements

We appreciate the open communication with Dr. Hugo Naya and the helpful comments from Adam Eyre-Walker.

## Funding

This work was supported by the National Natural Science Foundation of China (31671321, 31421063, and 31371283).

## Availability of data and materials

The data generated and analysed during this study are included in the Additional files (Additional file 2).

## Authors’ contributions

D.K.N. conceived the study and wrote the manuscript. S.A. retrieved the data from online databases, matched the pairs, calculated the genomic GC content and the 16S rRNA identity, and performed the statistical tests. X.R.L. identified the orthologous genes and the repairing enzymes, calculated the GC content and nucleotide substitution rates at 4FDS. B.W.Z. identified the 16S rRNA genes. ZLC verified some of the results. All authors read and approved the final manuscript.

## Ethics approval and consent to participate

Not applicable.

## Competing interests

The authors declare that they have no competing interests.

